# Fibroblast alignment is governed by stiffness interfaces in hydrogels

**DOI:** 10.64898/2025.12.19.695439

**Authors:** R.J.B. Dijkstra, J.K. Burgess, X. Petridis, P.K. Sharma, M.C. Harmsen

## Abstract

Transitions in mechanical properties between different tissue scaffolds, such as extracellular matrices (ECMs), are common in the human body and influence cellular orientation. For example, in teeth, the abrupt transition between soft (∼5 kPa) dental pulp and stiff (>100 kPa) dentin directs the alignment of odontoblasts. In this study, hydrogels, with defined mechanical transitions were used to evaluate the phenotypic response of fibroblasts to either a soft/soft (3 kPa/3 kPa) or soft/stiff (3 kPa/200 kPa) interface. Near the soft/soft interface fibroblasts aligned in a more parallel fashion to the interface (11° angle to the interface), compared to the more perpendicular fashion (69° angle to the interface) observed at the soft-stiff interface. Notably, regions of alignment also showed an increased proportion of alpha smooth muscle actin (αSMA)-positive cells, indicating myofibroblast differentiation in response to the mechanical transition. These findings demonstrate that modifying a biomaterial by introducing an interface alters the bioactivity of responding cells solely through mechanical mechanisms, without any change to the cells’ biochemical environment.

Graphical abstract

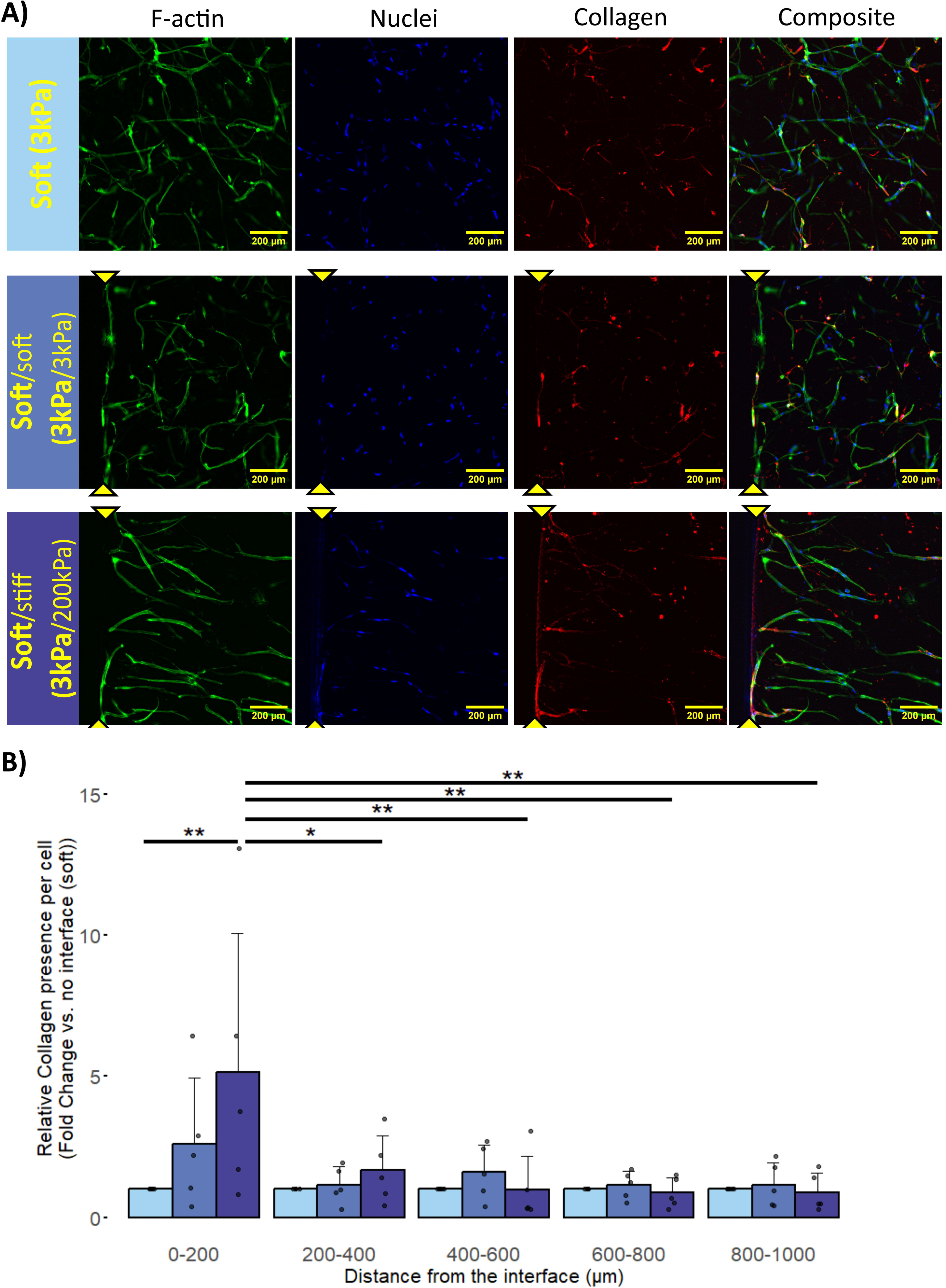

Schematic summary showing interface-induced fibroblast orientation and variety in phenotypic expression. On the bottom left the perpendicular orientation of fibroblast (skin tone) and myofibroblast (pink) on a soft/soft (3kPa/3kPa) GelMA hydrogel interface and on the right side the parallel orientation near a soft/stiff (3kPa/200kPa) GelMA hydrogel interface with greater collagen deposition, depicted by purple halo’s.

## Introduction

The extracellular matrix (ECM) is a three-dimensional network undergoing continuous remodeling of its structural components including proteoglycans, collagens, and elastin, which together with the tissue’s water content, collectively determine tissue stiffness (Olivares-Navarrete et al., 2017; Yi et al., 2022). The stiffness of ECM varies markedly across different tissues, ranging from 0.1-1 x10^3^ pascal (Pa) for brain or adipose tissue, 8-17 x10^3^ Pa for muscle to 25-40 x10^6^ Pa for osteoid matrices (Engler et al., 2006), supporting each tissue’s specific functional demands (Padhi & Nain, 2020). Variations in the mechanical properties of the ECM influence cellular phenotypes, affecting differentiation, migration, morphology, and orientation (D’Urso & Kurniawan, 2020; Sung et al., 2022; Yi et al., 2022).

Maintenance of normal tissue homeostasis depends on the reciprocal, balanced and often dynamic interaction between cells and the ECM, ensuring mechanical equilibrium which suppresses pathological alterations (Di et al., 2023; Handorf et al., 2015). This balance is sustained through continuous matrix remodeling, primarily mediated by mesenchymal cells via cell–matrix interactions (Bonnans et al., 2014).

Matrix remodeling is initiated through mechanotransduction mediated by surface mechanoreceptors such as integrins. These mechanoreceptors connect the ECM to the cytoskeleton through the assembly of focal adhesions, enabling cells to sense and respond to mechanical cues (Martino et al., 2018; Sun et al., 2016; Weinberg et al., 2017). Through these adhesions, cells exert contractile forces on the ECM, with the magnitude of these forces influencing adhesion size (Tan et al., 2003). In turn, the size and number of focal adhesions regulates cellular contractility, contributing to cell differentiation, migration, morphology, and orientation in response to mechanical stimuli (Berginski et al., 2011; Saraswathibhatla et al., 2023). However, cellular contractility is not the sole source of tensile forces within the ECM. Heterogeneities in the intrinsic material properties-particularly variations in macromolecular concentration and the number of interactions per molecule, also contribute to the buildup of internal tension (Hong & Wang, 2013). These intrinsic material properties also contribute to the ECM stiffness (Frantz et al., 2010), where abrupt transitions in stiffness (interfaces) can influence cellular behavior such as differentiation and orientation (Bonnans et al., 2014).

Abrupt transitions in stiffness between soft (dental pulp) and hard tissues (dentin) give rise to region specific cell organization and function. At these interfaces, where ECM regions of distinct stiffness meet, stress fields develop due to sharp mechanical discontinuities (Z. Zhou et al., 2017). The magnitude of these interfacial stresses depends on the degree of kinematic constraint between adjoining regions, potentially generating surface tension and localized mechanical gradients (Duo et al., 2025) . Despite growing evidence that ECM stiffness modulates cell behavior and alignment (Athirasala et al., 2017; Ha et al., 2020), the specific cellular responses to abrupt stiffness transitions between adjacent matrix domains remain insufficiently understood.

To address this knowledge gap in the present study, we employed gelatin methacrylate (GelMA) hydrogels to engineer interfaces between regions of distinct stiffness. GelMA was selected due to its tunable stiffness as well as its biocompatibility and ability to support cell viability and differentiation (Khayat et al., 2017; Kirsch et al., 2019; Pepelanova et al., 2018). In previous work, we showed that stress relaxation hardly varies between 5 and 15% (w/v) GelMA, resulting in viscoelastic stability (Martinez-Garcia et al., 2021). Combined with its ability to form networks through photo-crosslinking, GelMA provides an ideal platform for constructing matrices with abrupt stiffness transitions while incorporating seeded cells.

In this study we investigate how stiffness transitions influence cellular phenotype, and therewith if biomaterial mechanical modulation alone, under consistent biochemical conditions, is sufficient to provoke phenotypic cell responses.

## Materials and Methods

### GelMA synthesis

Methacrylated gelatin (GelMA) was synthesized according to the one-pot method described by Shirahama et al. (Shirahama et al., 2016). Briefly, 20 gr of type A porcine gelatine (Bloom strength 300, Merck KGaA, Darmstadt, Germany) was dissolved in 200 mL carbonate buffer (0.25 M). Subsequently, 2 mL of methacrylic anhydride (MAA; Merck KGaA) was added, and the mixture was stirred at 50°C for 2h. The solution was then dialyzed (14 kDa, Merck KGaA, Darmstadt, Germany) for 3-4d at 40°C, with water refreshments every 4 to 16h. Finally, the GelMA was lyophilized (Labconco Corporation, Kansas City, MO, USA) for 2-3d and stored at - 20°C.

### Degree of functionalisation

The 2,4,6-Trinitrobenzenesulfonic acid (TNBS, Merck KGaA, Darmstadt, Germany) assay was used to quantify the degree of functionalization (DOF) (Loessner et al., 2016). By taking non-functionalized gelatine (100%) and comparing it to functionalized GelMA with respect to the substituted lysine groups, the degree of functionalization can be calculated. Briefly, GelMA and unfunctionalized gelatin were separately dissolved in 0.1 M NaHCO_3_ (Merck KGaA, Darmstadt, Germany) to a final concentration of 500 μg/mL. After serial dilution (500 μg/mL, 250 μg/mL, 125 μg/mL, 62.5 μg/mL, 31.25 μg/mL and 0 μg/mL) TNBS solution (0.01%) was added in a 1:2 ratio. After 2h at 37°C in the dark, optical density (OD) at 335 nm was measured. The mean OD of triplicate measurements was plotted and the slopes of both the GelMA and unfunctionalized gelatine were determined. The DoF was calculated using equation 1;

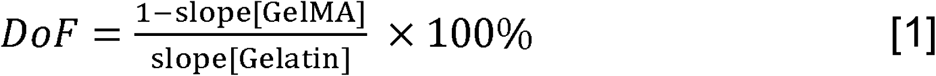

### Cell culture

Human lung fibroblasts (MRC5, ATCC CCL-171, Molsheim Cedex, France), negative for *Mycoplasma spp*, were cultured in high glucose Dulbecco’s modified Eagle medium (DMEM; Lonza, Walkersville, USA) containing 10% Fetal Bovine Serum (FBS; Sigma-Aldrich), 1% Penicillin–Streptomycin (Pen-Strep; Gibco^TM^, Paisley, Scotland) and 2 mM L-glutamine (BioWhittaker®, Verviers, Belgium) and cultured at 37°C, 5% CO_2_. Upon reaching 80–90% confluency, fibroblasts were detached using 0.5% trypsin-EDTA (Gibco™). Cells were then centrifuged at 300LJ×LJg for 5 min, resuspended in fresh medium, and counted using a NucleoCounter NC-200 (ChemoMetec, Allerød, Denmark).

### Stiffness interface preparation

To construct a hydrogel with a stiffness transition, an interface between soft (3LJkPa; 5–7% w/v GelMA) and stiff (200LJkPa; 15% w/v GelMA) regions was prepared. First, 100 μL of either the soft or stiff GelMA solution, mixed with 0.1% lithium phenyl-2,4,6-trimethylbenzoylphosphinate (LAP; Tokyo Chemical Industry UK Ltd), was dispensed into one half of an Eppendorf tube lid (Eppendorf, Netherlands), with the remaining space blocked by a spacer (Provil® novo Putty: soft set fast, Kulzer GmbH, Wasserburg, Germany). The solution was then photopolymerized using 405 nm light for 5 min, ensuring all available crosslinking sites were fully utilized. After removal of the spacer, the second hydrogel solution (soft or stiff), containing encapsulated cells at 1 × 10LJ cells/mL and supplemented with 0.1% LAP, was added adjacent to the first. UV-A photopolymerization (405 nm, 5 min) of the second adjacent part of the hydrogel ensured gelation and crosslinking, forming a covalently bonded interface (Fig 1A).

**Figure 1:**
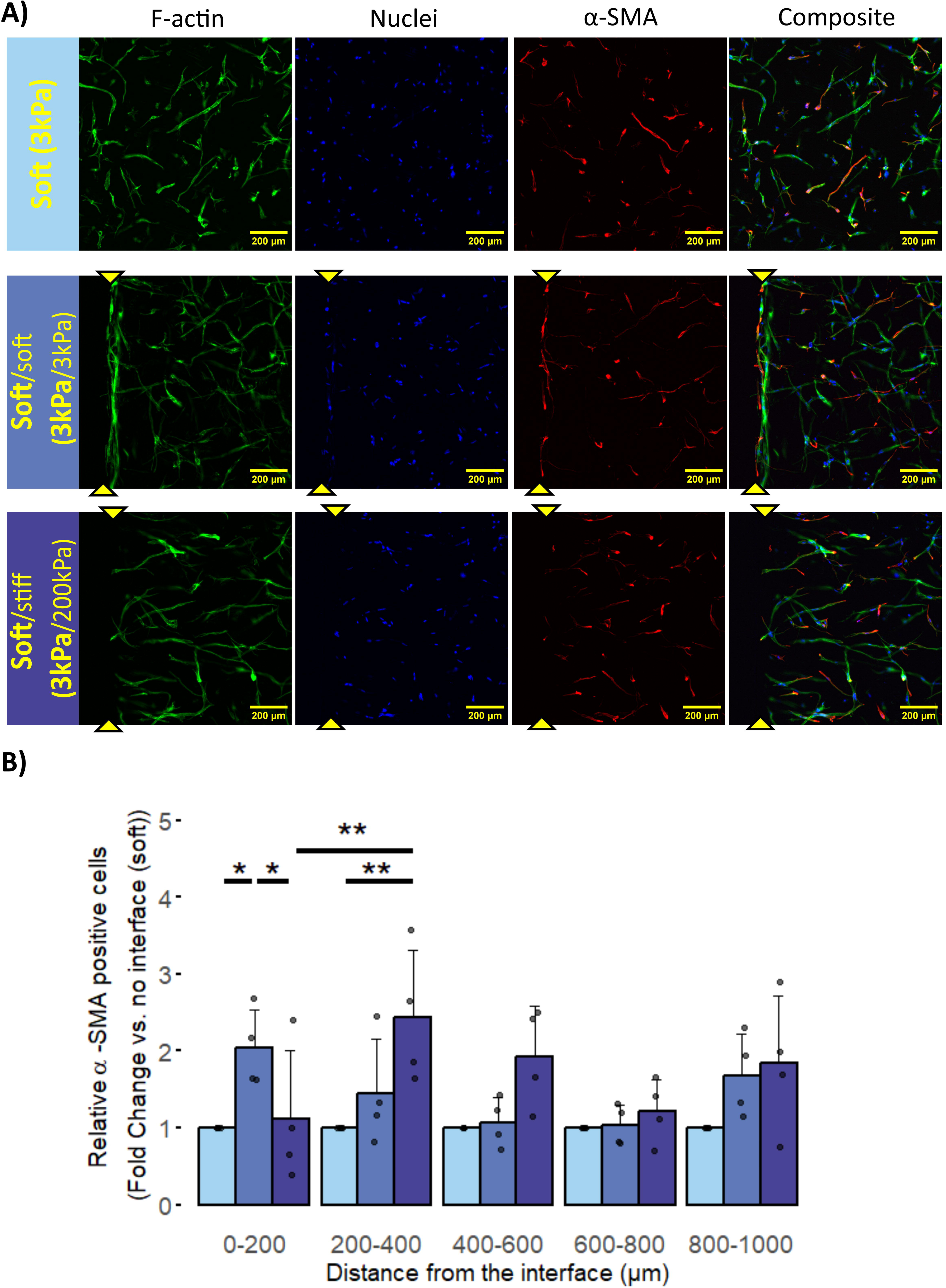
A) Schematic illustratio of the construction of a GelMA hydrogel interface composed of two regions with distinct stiffness: a soft domain (31kPa, light blue) and a stiff domain (2001kPa, purple). This interface represents an abrupt stiffness transition designed to study its influence on cellular phenotypic responses. B) Schematic of GelMA hydrogels with encapsulated cells (bold font). The top row shows stiff GelMA hydrogels (2001kPa, purple), and the bottom row shows soft GelMA hydrogels (31kPa, light blue). Encapsulated cells (light brown) with nuclei (dark blue) are depicted. The first column represents control hydrogels without an interface, while the second and third columns show constructs with interfaces between stiff and soft regions, with cells encapsulated. C) Photograph of a GelMA hydrogel with an interface. At the left side soft cell containing hydrogel (1 × 10⁶ fibroblasts/mL) and adjacent to it on the right cell free stiff hydrogel.

Control hydrogels were prepared by filling the entire lid with 200 μL GelMA (with or without encapsulated cells), either soft or stiff. The following conditions were analyzed:

Control groups: soft, (∼3kPa), and stiff (∼200kPa) hydrogels (each with and without cells); Experimental: **soft**/stiff (∼3kPa/∼200kPa), **soft**/soft (∼3kPa/∼3kPa), **stiff**/stiff (∼200kPa/∼200kPa), and **stiff**/soft (∼200kPa/∼3kPa)hydrogels (Fig 1B).

**Bold font** indicates where the cells were encapsulated within the interface hydrogels.

All hydrogel variations were cultured, free floating, in a 12 well cell culture plate (Corning Incorporated, Kennebunk, USA) in high glucose DMEM containing 10% FBS, 1% Pen-Strep and 2mM L-glutamine at 37°C, 5% CO_2_ for up to 14d.

### Mechanical properties characterization

Hydrogel stiffness was assessed using a low-load compression tester (LLCT) at days 0, 3, 7, and 14. A plunger (2.5 mmφ) compressed each hydrogel to 80% of its original height at a strain rate of 0.2 s⁻¹. Stiffness was calculated from the slope of the stress–strain curve. Stress relaxation at constant strain (ε = 0.2) was monitored over 100 seconds and expressed as the percentage of stress relaxation (%R)(Peterson et al., 2013).

The correlation between concentration (% GelMA) and stiffness (Pa) was assessed by measuring a calibration curve using GelMA concentrations of 5, 6, 7, 8, 10 and 15% (w/v). The correlation (equation 3) was then used to determine the needed GelMA concentration to obtain hydrogels with a stiffness of 3 and 200 kPa, respectively.

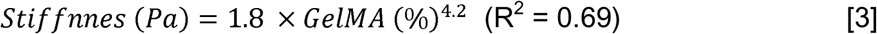

### Immunohistochemical staining

Cell morphology and orientation were assessed by fluorescent labeling of the cytoskeleton (Phalloidin-iFluor 488, Abcam) in conjunction with nuclei-staining with (4′,6-diamidino-2-phenylindole, DAPI; Sigma–Aldrich) to identify the cell count. At 0, 3, 7 and 14d of incubation, cell-laden hydrogels were fixed overnight in 4% paraformaldehyde, then rinsed three times with Dulbecco’s phosphate-buffered saline (DPBS; BioWhittaker®, Walkersville, MD, USA) for five min. Cells in the hydrogels were permeabilized with 0.1% Triton X-100 in DPBS for 10 min. Blocking was performed by incubating with 1% bovine serum albumin (BSA) and 5% goat serum. Primary antibody (Ki-67 (1:200) or COL1A1 (1:200) or αSMA (1:200)) was diluted in DPBS containing 1% BSA and 5% goat serum, followed by overnight incubation at 4°C. Hydrogels were rinsed using DPBS supplemented with 0.05% Tween. The secondary antibody (Alexa Fluor 568, Abcam) was diluted (1:500) in DPBS containing phalloidin-iFluor 488 (1:1000) and DAPI (1:5000). After 60 min incubation at room temperature hydrogels were rinsed with DPBS supplemented with 0.05% Tween. Image stacks over the interface in the middle of the sample, composed out of five images over a depth of 200 µm, were recorded using a Thunder Imaging system (Leica microsystems, Amsterdam, The Netherlands). Image stacks of 5 slices with an interslice spacing of 50 µm, covering a total depth of 200 µm were processed in FIJI (ImageJ v1.54f) (Schindelin et al., 2012) and projected to 2D images. The projected volume was considered the compacted area in the 2D image.

### Cell morphology and Orientation

In order to analyze the cell orientation and morphology from the immunofluorescent staining, object separation was accomplished using morphological segmentation in FIJI (ImageJ v1.54f) . Image input was set to Object with a gradient radius of 2 and watershed lines were enabled, creating an object separation mask. Subsequently the separation mask was applied to the fluorescent image, creating an image with separate cells.

The cell morphology was determined by the length-width ratio derived from each phalloidin-labelled cell cytoskeleton in an image. In FIJI (ImageJ v1.54f) the length on the major axis and the width on the minor axis of each cell were measured and recorded and equation 2 was used to calculate the Aspect Ratio (AR) for each cell. When the aspect ratio exceeded a value of two, the cell was considered elongated.

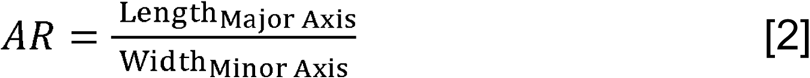

Subsequently, the percentage of elongated cells in each of hydrogel condition was determined by taking the number of elongated cells and dividing it by the total number of cells found.

To assess the orientation the angle between the cell’s major axis and the interface was determined. The cell center of mass was used as a reference point to measure its distance from the stiffness interface. Cell location was binned in 200 μm intervals over a range of 0-1000 μm. For interface free hydrogels a random point was used as reference point for the distance binning. For each distance bin, the cell angles were recorded where all the angles >90° were reflected to a 0–90° scale by subtracting the angle from 180°.

### Matrix remodeling and Cell differentiation

As a measure of matrix remodeling the degree of collagen deposition, visualized by immunofluorescent staining (anti-COL1A1) was quantified as the fold change in collagen area (um^2^), normalized for the number of cells, between interfaces containing hydrogels **soft**/soft and **soft**/stiff and the **soft** (3kPa) hydrogel without an interface. The relative collagen area (μm^2^) per cell was determined within each of the previously mentioned distance bins (200 μm intervals (0-1,000 μm). In short, the collagen area (μm^2^) in each image was measured and divided by the number of nuclei found in the corresponding bin. The collagen area (μm^2^) per cell found in the **soft** (3kPa) hydrogel without an interface was considered as control and set to 1.

Subsequently, the relative collagen per cell was calculated in each distance bin (200 μm intervals (0-1,000 μm) for the interface containing hydrogels **soft**/soft (3kPa/3kPa) and **soft**/stiff (3kPa/200kPa) and presented as the fold change in relative collagen deposition.

The percentage of αSMA-positive cells, quantified similarly as above, was used to assess fibroblast to myofibroblast differentiation. Immunofluorescent staining (anti-αSMA) was used to detect αSMA positive areas, larger than 50 um^2^ (diameter of ∼8 μm). Colocalization of the actin-skeleton plus nuclei and the αSMA positive areas gave rise to the number of αSMA positive cells per hydrogel type. The number of αSMA positive cells found in the **soft** (3kPa) hydrogel without an interface was considered as control and set to 1. Subsequently, the number of αSMA positive cells was calculated in each distance bin (200 μm intervals (0-1,000 μm) for the interface containing hydrogels **soft**/soft (3kPa/3kPa) and **soft**/stiff (3kPa/200kPa) and presented as the fold change in αSMA positive cells and interpreted as a measure for fibroblast to myofibroblast differentiation.

### Statistical analyzes

Data from at least five unique samples per combination were analyzed in R (3.6.0+) using Rstudio (Version: 2024.09.1+394, released: 2024-11-04). Data normality was assessed with Q-Q plots and Shapiro-wilk test. Subsequently, normally distributed data were analyzed using two-way ANOVA followed by Tukey post-hoc test. Non-normally distributed data were analyzed using Kruskal-Wallis test followed by Wilcoxon’s or Dunn’s post-hoc test. Statistical significance was set at p ≤ 0.05.

## Results

### Fabrication of stiffness-defined hydrogels

To assess the impact of abrupt stiffness transitions within a matrix, the degree of functionalization (DoF) of the GelMA batches was quantified to ensure consistent mechanical properties. The measured slopes for each of the three batches (6 replicates per batch) were 9.9LJ×LJ10⁻LJLJ±LJ2.1LJ×LJ10⁻LJ (OD /C (µg/mL), Batch 1), 10.6LJ×LJ10⁻LJLJ±LJ2.4LJ×LJ10⁻LJ (OD /C (µg/mL)), Batch 2), and 6.9LJ×LJ10⁻LJLJ±LJ2.1LJ×LJ10⁻LJ (OD /C (µg/mL)), Batch 3) (Supplementary 1A). GelMA used in all hydrogel samples exhibited an average DoF of 68.1LJ±LJ4.5% (Batch 1), 65.5LJ±LJ7.5% (Batch 2), and 70.7LJ±LJ5.3% (Supplementary 1B).

Initial stiffness values of 3LJkPa and 200LJkPa were selected for hydrogel fabrication. Using our established model, GelMA concentrations of approximately 5.8% and 15.9% were extrapolated to achieve stiffness values of 3LJkPa and 200LJkPa, respectively.

### Fibroblast-mediated modulation of hydrogel stiffness

To assess the impact of cellular activity on matrix mechanics, fibroblasts were encapsulated in both soft and stiff GelMA hydrogels and monitored for 14 days. In soft hydrogels, initially, encapsulating fibroblasts increased the stiffness from 2.9 kPa without fibroblasts to 4.7 kPa with fibroblasts (p=0.01). Over time the presence of fibroblasts reduce the matrix stiffness (day 0 4.7 kPa, day 3 3.7 kPa, day 7 3.4 kPa and day 14 2.3 kPa, (H(3)LJ=LJ9.4, pLJ=0.02)), however in the absence of fibroblasts the stiffness did not change (Table 1). After 14 days the stiffness of the matrix in soft hydrogels with and without fibroblasts was equivalent.

In contrast, fibroblasts encapsulated in stiff hydrogels reduced matrix stiffness from 229LJkPa without fibroblasts to 138LJkPa with fibroblasts (p=0.01) at day 0. After the initial stiffness reduction no further change in stiffness was observed in the presence of fibroblasts. At day 0 the stiffness was measured to be 138 kPa, decreasing to 100 on day 3, 109 kPa at day 7 and after 14 days the stiffness was found to be 107 kPa (H(3)=2.3, p=0.51) (Table 1). In the absence of fibroblasts there was a slight decrease in stiffness of the stiff hydrogel. The initial stiffness at day 0 of 229 kPa lowered to 194 kPa on day 3, to 190 kPa on day 7 and ended up at 173 kPa at day 14 (H(3)=10.7, p=0.01).To evaluate the impact of fibroblasts on the viscoelastic properties of GelMA hydrogels, stress relaxation was measured over the 14-day culture period. Initially, encapsulating fibroblasts in the soft hydrogel did not change the stress relaxation at day 0 (5.9% without, 5.6% with fibroblasts). In the presence of fibroblasts, after day 14, an increase in stress relaxation was observed in the soft hydrogel, respectively 6.9 % vs. 8.1 % (p=0.02) (Table 2). Whereas the stress relaxation in the soft hydrogel did not change over 14 days when no fibroblasts were present (H(3)=2.8, p=0.42)(Table 2).

In stiff hydrogels, a similar pattern to that observed for stiffness was present. The stress relaxation dropped from 6.1% without to 3.9% with fibroblasts (Table 2). Over the 14 day culture period the change in stress relaxation remained the same.

### Stiffness interfaces direct fibroblast morphology and alignment

To investigate how stiffness interfaces influence fibroblast morphology and alignment, cells were encapsulated in various hydrogel combinations and cultured for 14 days (Fig. 2 (day 14), supplement 2 and 3 (day 3 and day 7). Elongation of fibroblasts, represented by a length-width ratio greater than 2, was increased in soft hydrogels, whereas in stiff hydrogels elongation was absent (supplement 4). The percentage of elongated fibroblasts increased in the interface free soft hydrogel from 31.2 ± 21.5% at day 3, to 38.8 ± 3.0 at day 7, to 59.5 ± 16.2% at day 14 (Fig. 4). The same trend was found in soft hydrogels where an interface was present (where **bold** font indicates the fibroblast containing hydrogel). For **soft**/soft hydrogels the percentage elongated fibroblasts increased from 26.1 ± 13.1% at day 3 to 49.4 ± 2.9% at day 7 ending up at 60.7 ± 12.3% at day 14 (Fig 3). In the hydrogel with a **soft**/stiff interface the increase in fibroblasts elongation was most prominent, with 18.9 ± 5.0% at day 3, compared to 48.0 ± 4.9% at day 7 and 67.6 ± 12.3% at day 14 (Fig. 3). While the percentage of elongated fibroblasts remained 10% or lower for all hydrogel samples in which the fibroblasts were encapsulated in the stiff part (**stiff**, **stiff**/soft and **stiff**/stiff).

**Figure 2.**
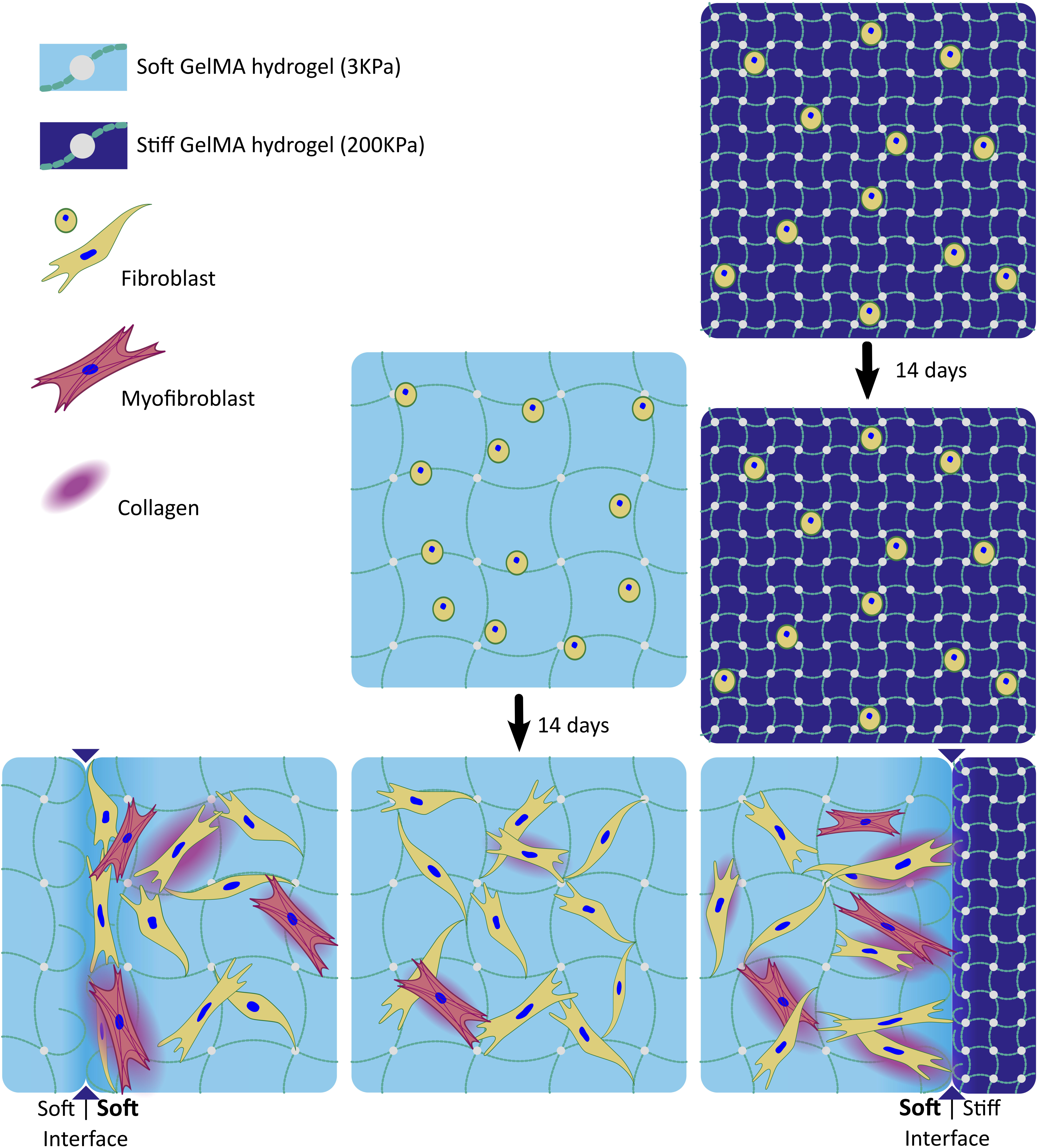
: Stiffness transition driven fibroblast morphology and orientation. Fibroblast morphological and orientation differences after 14 days, influenced by the interface between GelMA hydrogels (indicated by the yellow arrows) composed of either soft (3kPa) or stiff (200kPa) hydrogel. The cell-containing hydrogel is indicated in bold font (1x10 fibroblasts/ml). The cytoskeleton (Phalloidin) is shown in green and nuclei (DAPI) in blue. The scale bar represents 200 μm. Images are representative of three experimental repeats.

**Figure 3.**
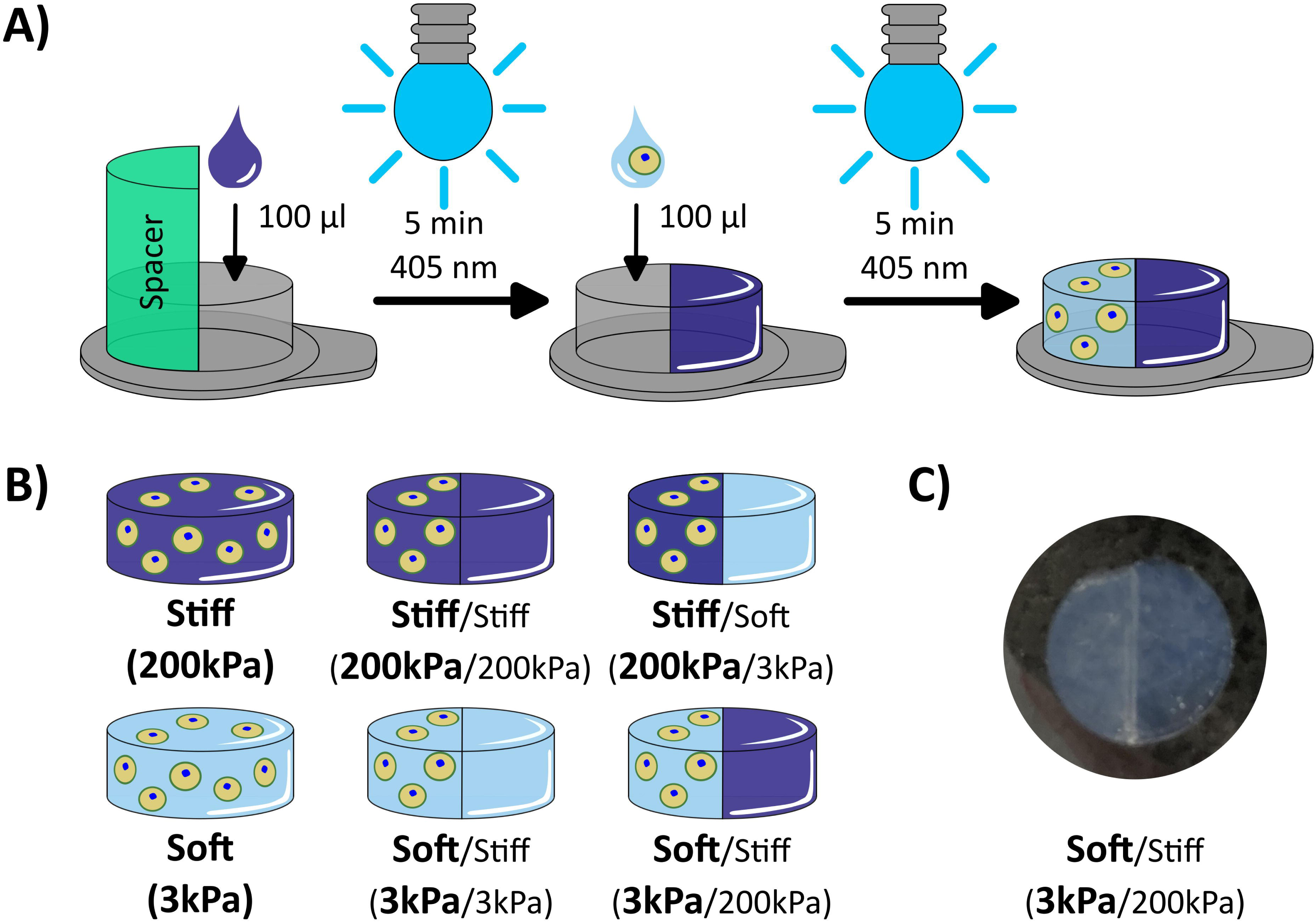
: Stiffness interfaces direct fibroblast morphology. Hydrogel stiffness dependent percentage elongated fibroblast after culture for 3 (light pink), 7 (purple), and 14 (dark burgundy) days. Bold font indicates the cell-containing hydrogel with a seeding density of 1x10^6^ fibroblasts/ml. The mean percentage elongated fibroblasts was calculated from a minimum of 750 cells per individual soft, soft/soft, soft/stiff, stiff, stiff/soft and stiff/stiff hydrogels, at least three hydrogels per time point were measured (black dots). ANOVA with Tukey post-hoc analysis was used to assess differences between the groups. (* p<0.05, ** p<0.01 and *** p<0.001).

**Figure 4.**
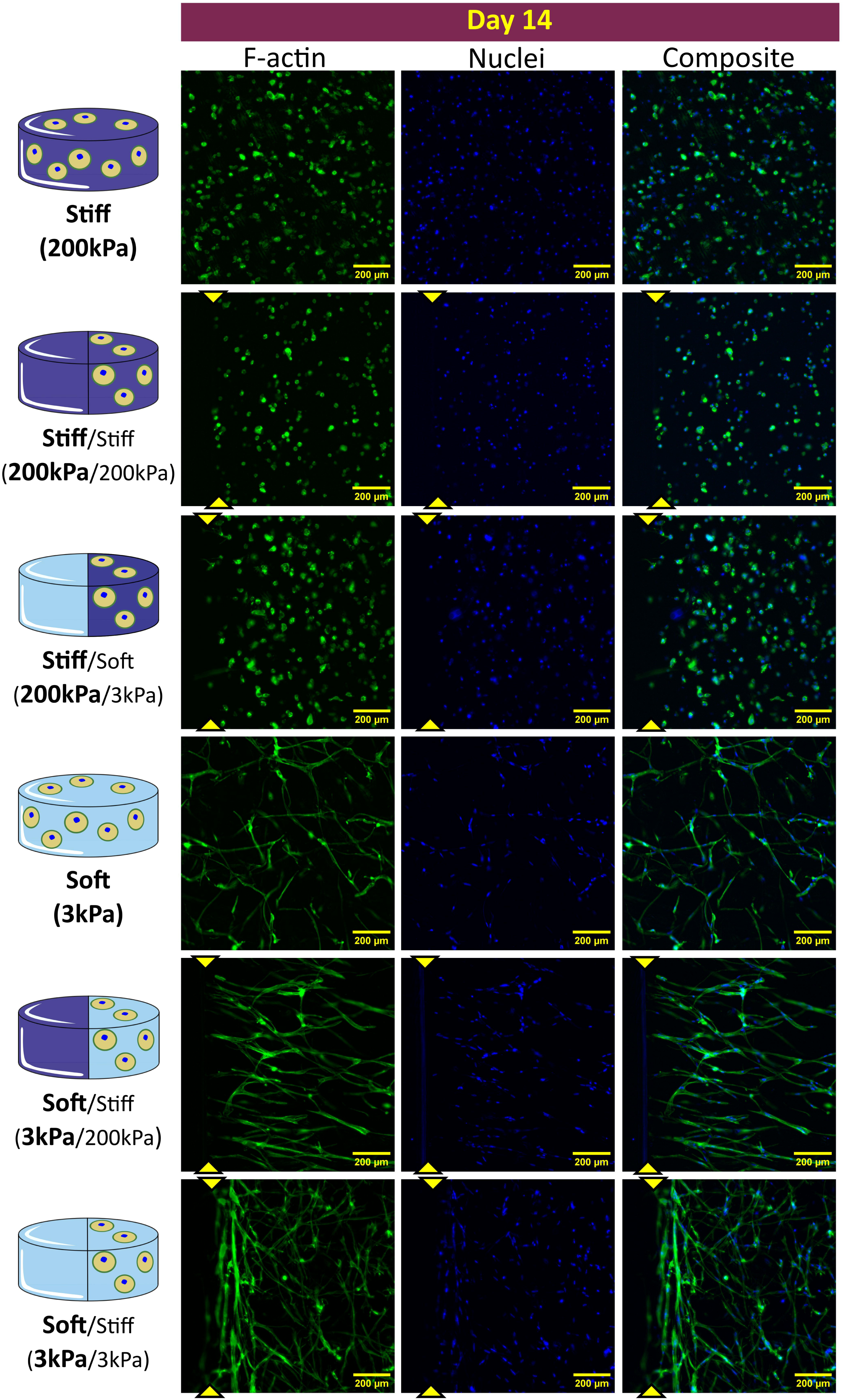
: Stiffness interfaces direct fibroblast alignment. A) The distribution in cell angle of fibroblasts (°) relative to the interface within GelMA hydrogel at day 14 of culture. The fraction of cell angles is plotted for soft (light blue), soft/soft (lavender), and soft/stiff (purple). B) Median cell angle relative to the interface derived from at least 30 fibroblasts per distance in at day 14 of culture for soft (light blue), soft/soft (lavender), and soft/stiff (purple). Statistical differences were assessed using Kruskal-Wallis with Dunn’s post hoc test (*p < 0.05, **p < 0.01, ***p < 0.001). From left to right, the distance from the interface is given in bin ranges of 200 μm, up to 1000 μm distance from the interface.

In interface free soft hydrogels, elongated fibroblasts exhibited random orientation, with a median orientation angle of approximately 45° (range: 40°–58°; IQR: 17°–76°). However, in soft hydrogel, fibroblasts located in the range of 0-200 μm from the interface with soft hydrogel aligned their long axis parallel to the interface, with median angles of 17° (IQR: 5°–49°) at day 3, 14° (IQR: 6°–32°) at day 7, and 11° (IQR: 5°–31°) at day 14 (Fig. 4 (day 14), supplement 3 (day 3) and 4 (day 7)). Beyond 200 μm from the interface, the cell orientation became random again, with median angles ranging from 34° to 50° across all time points (H(2)=2.9, p=0.24).

In contrast, fibroblasts in soft hydrogel near an interface with a stiff hydrogel aligned their long axis in a more perpendicular fashion to the interface. This alignment was most pronounced at distances of 200–400 μm (68°, IQR: 52°–81° at day 7; 64°, IQR: 48°–81° at day 14) and 400–600 μm (69°, IQR: 47°–83° at day 7; 62°, IQR: 34°–75° at day 14) (Fig. 5). At distances beyond 600 μm from the interface, fibroblast orientation remained random throughout the 14-day culture period (Fig. 4).

**Figure 5:**
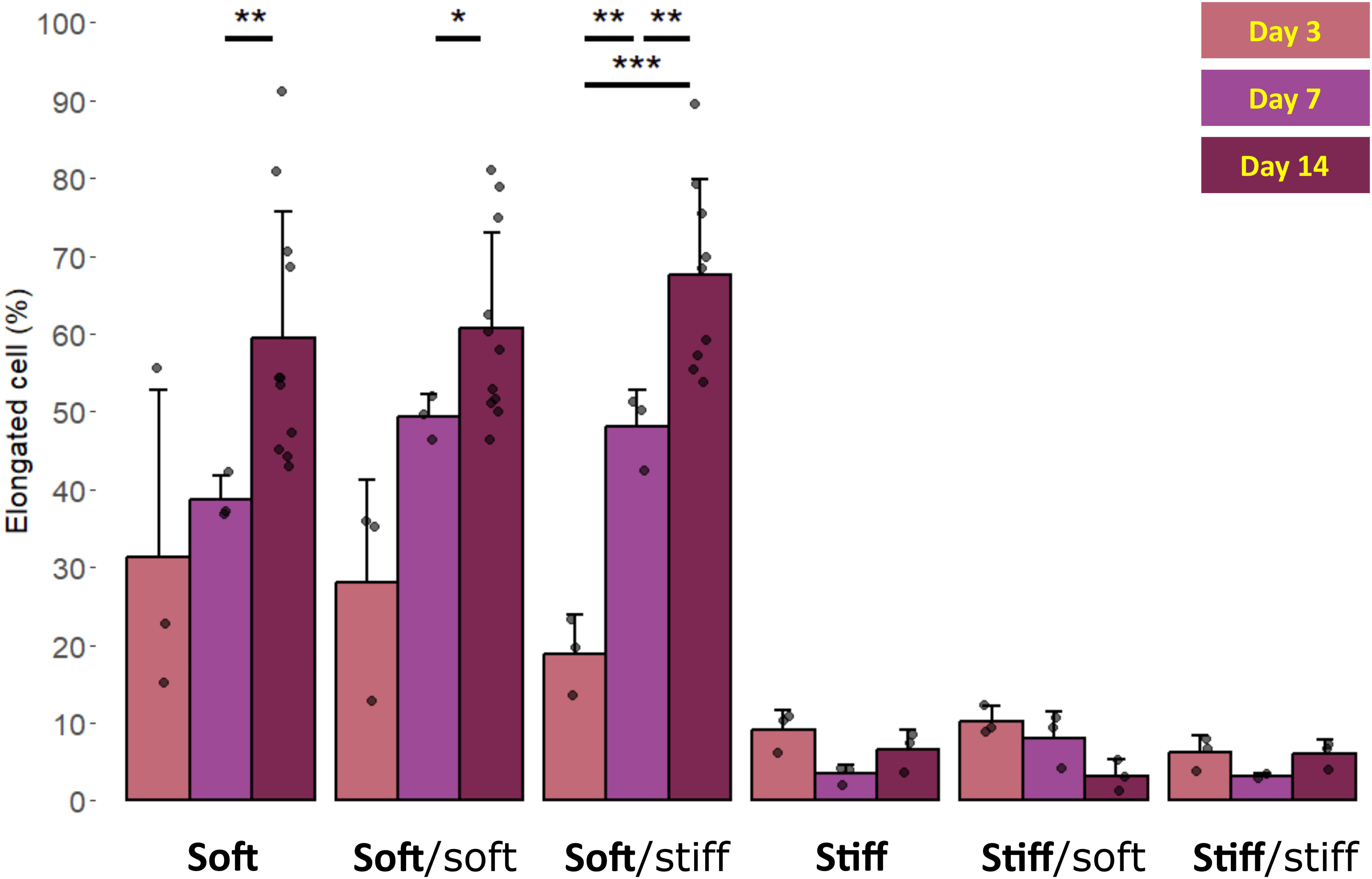
Stiffness interface influenced fibroblast distribution Mean ± standard deviation of the nuclei density (nuclei/mm²) at various the distances from the interface as a function of time. The distance to the interface is arranged in bins with a step size of 200 μm, ranging from 0 to 1000 μm from the interface. Within each distance bin, the nuclei density (n > 50 cell) is shown for day 3 (light pink), 7 (purple), and 14 (dark burgundy) of culture. From top to bottom, the different abrupt interface transitions are depicted: soft (light blue), soft/soft (lavender) and soft/stiff (purple). Bold font indicates the cell-containing hydrogel with a seeding density of 1×10⁶ fibroblasts/ml.

### Fibroblast density as a function of distance to interface

To assess the influence of stiffness interfaces on cell distribution, fibroblast density was quantified as a function of distance from the interface across all hydrogel combinations over the 14-day culture period. Overall, cell density remained relatively stable and did not exhibit a consistent temporal or spatial response with respect to the interface, regardless of stiffness pairing. The average nuclear density in **soft** hydrogels was 83LJ±LJ37LJnuclei/mm², the presence of an interface did not influence the overall cell density over the entire culture period (Fig 5).

### Interface proximity-dependent collagen presence

As a measure of matrix remodeling, presence of collagen per cell was quantified in relation to the distance from the interface within each hydrogel condition. The amount of collagen per cell within 0-200 µm from the **soft**/stiff interface increased by 5.1LJ±LJ4.9-fold (*p*LJ<LJ0.01), in comparison to without an interface in a soft hydrogel.

Fibroblasts at the **soft**/soft interface did not alter the collagen amount present in the same region (2.6LJ±LJ2.4%, *p*LJ=LJ0.9).

Beyond 200 μm from the interface (i.e., within the 200–1,000 μm region), collagen presence per cell was comparable across all hydrogel types. However, values remained lower than those observed in the 0–200 μm region of the soft/stiff condition, indicating a localized enhancement of matrix remodeling near the stiffness interface (Fig. 6).

**Figure 6:**
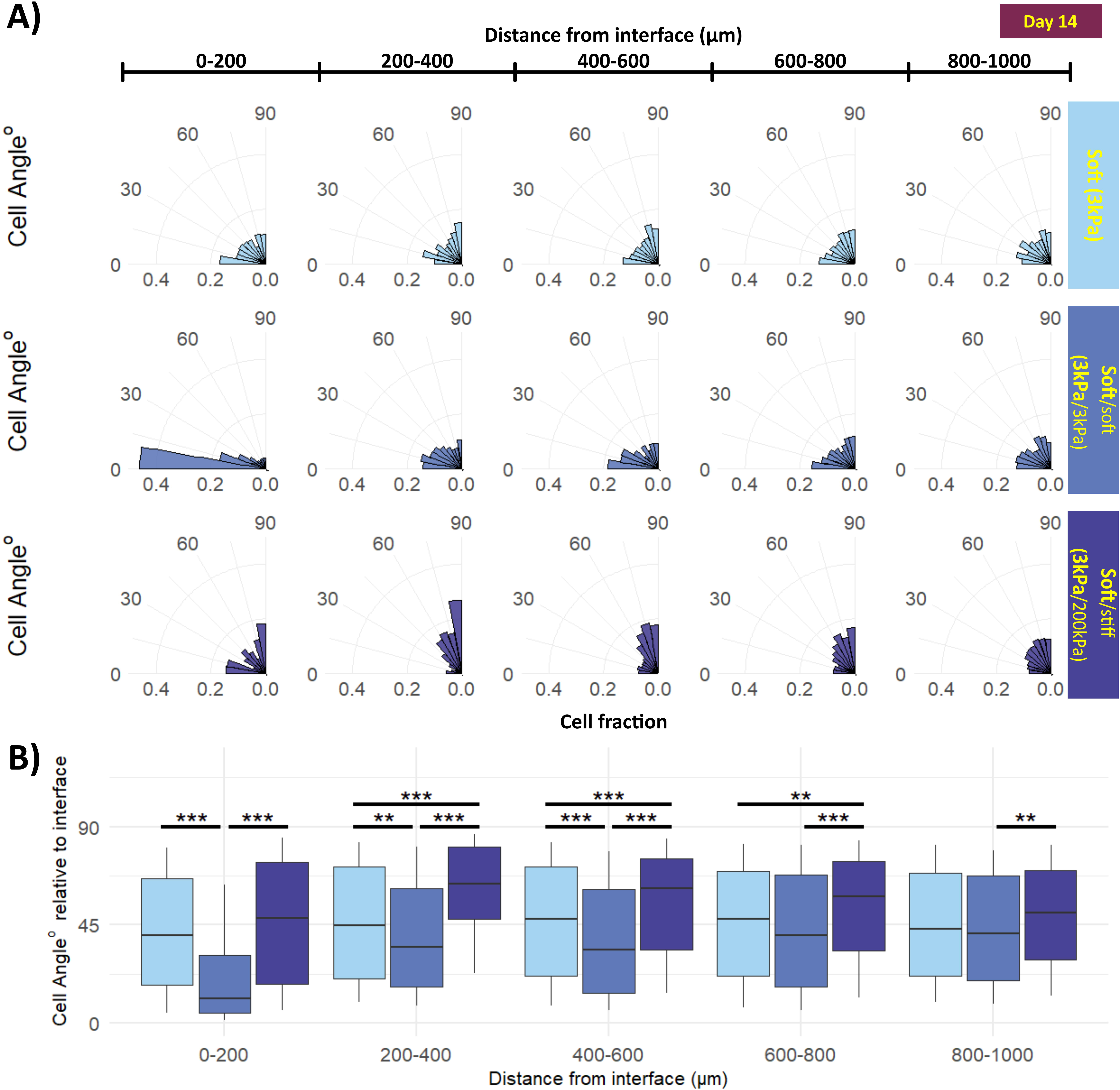
Distance to interface dependent collagen presence A) Immunofluorescent staining of fibroblast in soft (top), soft/soft (middle) and soft/stiff (bottom) hydrogel. F-actin (phalloidin stain (green), first column) shows the cell morphology and orientation, nuclei (DAPI stain (blue)) are shown in the second column. In the third column the presence of collagen (anti-COL1A1, red) is visualized and in the fourth column the composite image of the three stains is displayed. The interface is highlighted by the yellow pointers. The scale bar represents 200 µm. B) The fold change in relative collagen presence per fibroblast is shown as a function of distance from the hydrogel interface after 14 days of culture (n = 5, black dots). Distances (µm) are binned in 200-µm increments from the interface. Hydrogels with different stiffness configurations are shown soft (light blue), soft/soft (lavender), and soft/stiff (purple). Bold font denotes cell-containing hydrogels seeded at 1 × 10⁶ fibroblasts/mL. Statistical differences were assessed using two-way ANOVA with Tukey’s post hoc test (*p < 0.05, **p < 0.01, ***p < 0.001).

**Figure 7:**
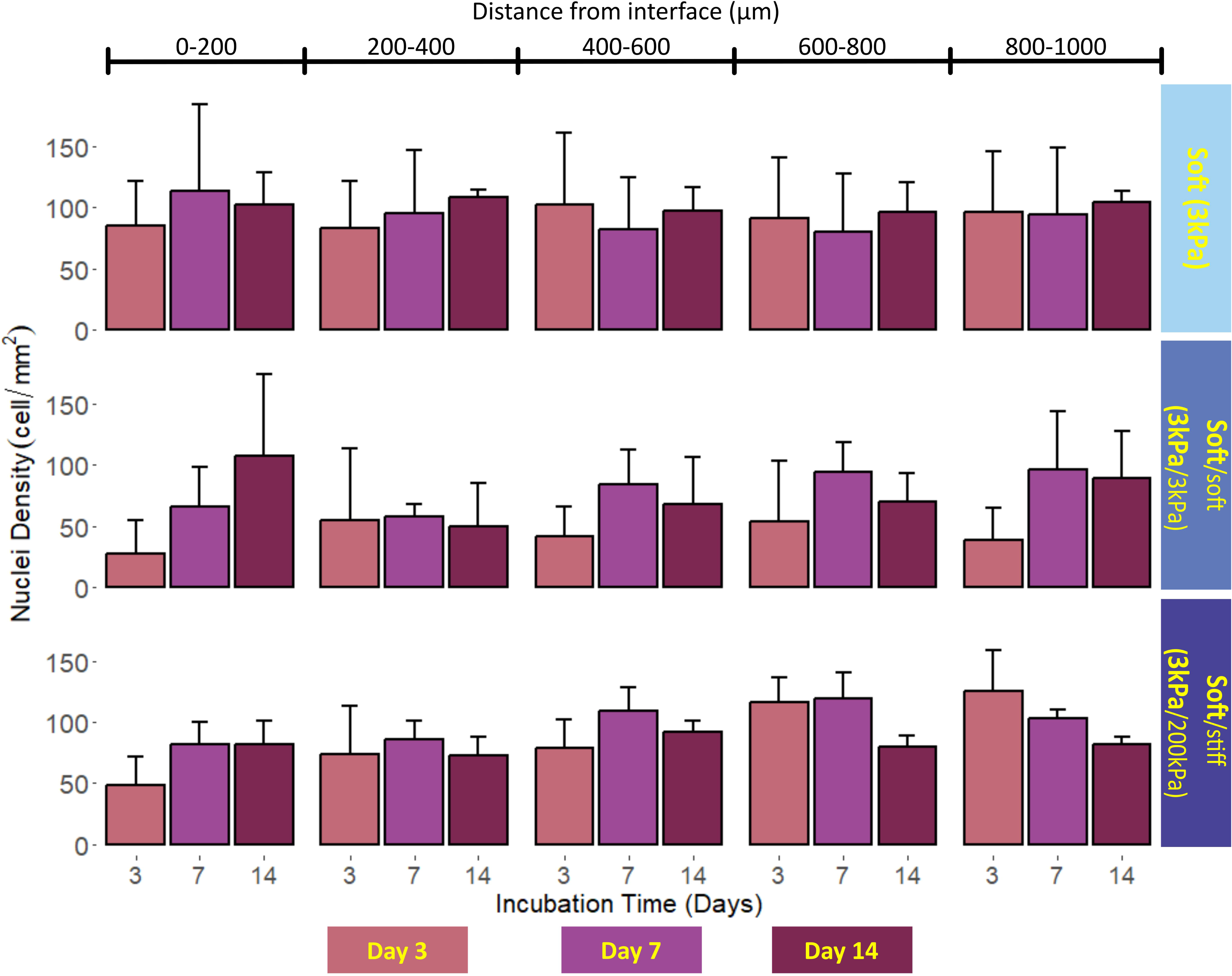
Distance to interface dependent fibroblast to myofibroblast differentiation. A) Immunofluorescent staining of fibroblast in soft (top), soft/soft (middle) and soft/stiff (bottom) hydrogel. F-actin (phalloidin stain (green), first column) shows the cell morphology and orientation, nuclei (DAPI stain (blue)) are shown in the second column. In the third column α-SMA presence (anti-α-SMA, red) is visualized and in the fourth column the composite image of the three stains is displayed. The interface is highlighted by the yellow pointers. The scale bar represents 200 µm. B) The fold change in α-SMA positive fibroblasts is shown as a function of distance from the hydrogel interface after 14 days of culture (n = 5, black dots). Distances (µm) are binned in 200-µm increments from the interface. Hydrogels with different stiffness configurations are shown soft (light blue), soft/soft (lavender), and soft/stiff (purple). Bold font denotes cell-containing hydrogels seeded at 1 × 10⁶ fibroblasts/mL. Statistical differences were assessed using two-way ANOVA with Tukey’s post hoc test (*p < 0.05, **p < 0.01, ***p < 0.001)

### Interface proximity-dependent myofibroblast differentiation

Fibroblast-to-myofibroblast differentiation (represented by αSMA positive cells) was assessed by quantifying αSMA expression and expressing the results as fold change relative to fibroblasts cultured in soft hydrogels without an interface. At a distance of 0-200 μm from the **soft**/soft interface, the number of αSMA positive fibroblasts was 2.0LJ±LJ0.5 times more in comparison to interface free soft hydrogel (p=0.05). And at the same distance from the **soft**/stiff interface, the number of αSMA positive fibroblasts, was 1.1LJ±LJ0.9 times more than in the **soft** hydrogel without an interface (p=0.02). The number of αSMA positive fibroblasts at a distance of 200-400 µm from the **soft**/stiff interface was 2.4LJ±LJ0.9 times more than in the same region for interface free **soft** hydrogel (*p*LJ<LJ0.01). Furthermore, αSMA positive fibroblasts were more abundant in the 200-400 µm region from the **soft**/stiff interface if compared to the 0-200 µm region in the same hydrogel (2.4LJ±LJ0.9 vs. 1.4LJ±LJ0.7 (p=0.001). Beyond 400 μm from the interface, fibroblast-to myofibroblasts gradually decreases with distance over the 14-day culture period.

## Discussion

Transitions in extracellular matrix (ECM) stiffness play a pivotal role in development, disease progression, and tissue engineering applications (Courbot & Elosegui-Artola, 2025). In this study, we investigated how abrupt stiffness transitions within GelMA hydrogels influence fibroblast behavior, including morphology, orientation, matrix remodeling, and differentiation.

Fibroblasts responded to stiffness transitions in distinct ways:

1. **Cell alignment** was dependent on the nature of the stiffness interface, with directional orientation emerging near **soft**/soft and **soft**/stiff interfaces.
2. **Matrix remodeling**, as measured by collagen presence per cell, was influenced by proximity to the interface in both the **soft**/soft and **soft**/stiff hydrogels.
3. **Differentiation to myofibroblast** was influenced by the presence of an interface and the proximity to it, with localized increases observed particularly in **soft**/stiff hydrogels.

In 3D matrices, cells are embedded within a surrounding structural network that can impose spatial constraints. When the matrix is too stiff, this network may act as a physical barrier, limiting cellular spreading and elongation, as previously described by Zhong et al. (Zhong et al., 2020), and reflect in our current findings on limited fibroblast spreading in the 3D cultures in stiff (200 kPa) GelMA hydrogels.

The ability of fibroblasts to elongate within a 3D GelMA hydrogel enabled the investigation of cellular alignment and motility in three dimensions. Distinct orientation patterns emerged at stiffness interfaces: fibroblasts aligned in a more perpendicular fashion to **soft**/stiff interfaces, whereas they adopted a more parallel fashion in alignment occurred at **soft**/soft interfaces. These patterns were spatially restricted, with perpendicular alignment most prominent at 200–600 μm from soft/stiff interfaces and parallel alignment confined to within 0–200 μm of soft/soft interfaces. This stiffness transition induced alignment of fibroblasts can be observed, for instance, in teeth. Where the soft dental pulp (∼5LJkPa) is encased by the much stiffer dentin (>100LJkPa), with odontoblasts aligning perpendicularly to the interface (Ronan et al., 2024). In addition to mechanical cues, chemical gradients (e.g., calcium) and physical gradients (e.g., oxygen) can also modulate cellular behavior.

Interestingly, despite pronounced changes in cell alignment and phenotype, fibroblast migration towards or across interfaces was not observed. This contrasts with numerous 2D studies where alignment and migration are closely linked (Ozcelikkale et al., 2017).

Non-random alignment, resulting from an active realignment in regions close to both the interface types, correlated with myofibroblastic differentiation, indicating a shift toward a motile and contractile phenotype characterized by αSMA expression. Notably, the spatial pattern of differentiation depended on the stiffness transition: in soft–stiff hydrogels, differentiation peaked 200–400 μm from the interface, whereas in soft–soft hydrogels, it was concentrated within 0–200 μm of the interface.

Collagen presence remained independent of myofibroblast differentiation. The highest collagen amount was observed near **soft**/stiff interfaces, despite a relatively low fraction of myofibroblasts in this region. This suggests that sharp stiffness transitions exert a profound influence on active matrix remodeling, likely driven by localized mechanical stress gradients (Kendall & Feghali-Bostwick, 2014). A biological example illustrating the functional relevance of increased collagen deposition at an abrupt stiffness transition can be found in teeth. Odontoblasts, which are aligned perpendicularly at the interface between the pulp (low stiffness) and dentin (high stiffness), exhibit specialized functionality in collagen secretion and mineralization, enabling the formation of new dentin (Linde & Goldberg, 1993).

The interplay between matrix stiffness transitions and fibroblast-mediated collagen presence indicates that these cells actively modulate local matrix architecture. Cells located near **soft**/stiff interface were associated with more collagen, consistent with previous findings that elevated matrix tension and stiffness regulate ECM production (Blache et al., 2020; Kolpakov et al., 1995). This interpretation is further supported by the absence of significant collagen accumulation at **soft**/soft interfaces.

Differentiation of fibroblasts into myofibroblasts occurred at both interface types, while in the **soft**/stiff matrix transition an region specific increase in collagen amount was observed, indicating that mechanical cues alone, without alterations in exogenous biochemical signals, can drive cell fate (Hussien et al., 2021; Kollmannsberger et al., 2018). These findings reinforce the concept that mechanical gradients within biomaterials act as potent regulators of cell phenotype.

A likely explanation for the observed influence of abrupt stiffness transitions on cell behavior is the generation of interfacial stresses arising from disparities in crosslinking density and molecular structure between adjacent hydrogels. These stresses propagate from the interface into the interior of the hydrogel, modulating focal adhesion dynamics and downstream actin cytoskeleton signaling (Greiner et al., 2013; Hong & Wang, 2013; Lee et al., 2018; Saraswathibhatla et al., 2023). It is intriguing to speculate that in **soft**/soft transition, the similar structures on either side of the interface would produce weak, short-ranged forces, resulting in parallel cell orientation. Conversely, the larger stiffness differential in **soft**/stiff interface is likely to generate long-ranged, intense forces that drive perpendicular alignment. Similar tension-induced cell reorientation has been reported in cyclic strain and shear stress models (Wen et al., 2009; Zhao et al., 2024). Unidirectional stress fields extend from soft/stiff boundaries under finite deformation (Hong & Wang, 2013), while biaxial deformations amplify both depth and directionality of these stresses (Y. Zhou & Jin, 2023). Notably, we did not observe migration across any interface, supporting the notion that high-density interfacial zones inhibit cell penetration unless a gradual stiffness gradient is present (Simona et al., 2015).

## Conclusion

Taken together, our findings emphasize the importance of stiffness transitions as a biophysical cue for directing fibroblast orientation and morphology. Our findings highlight the potential for leveraging mechanical interfaces to pattern tissue organization and induce cell specialization in engineered constructs without relying on exogenous growth factors. By precisely tuning stiffness at defined locations, biomaterials can be designed to instruct localized cell behavior, advancing regenerative strategies for complex tissues. Furthermore, our data demonstrate that mechanical signaling profoundly influences fibroblast phenotype, indicating that the bioactivity of a biomaterial can be modulated solely through mechanical design.

However, the intricate interplay between mechanotransduction and biochemical signaling within native tissue environments remains to be fully elucidated.

## Acknowledgments

The authors would like to thank Hans Kaper of the University Medical Center Groningen, department of Biomaterials and Biomedical Technology for his support in producing GelMA. This work was supported by Stichting De Cock - Hadders Foundation.

## Author contributions

Conceptualization: R.J.B.D., J.K.B., X.P., P.K.S., M.C.H. Data curation: R.J.B.D.

Formal analysis: R.J.B.D. Funding acquisition: R.J.B.D. Investigation: R.J.B.D.

Methodology: R.J.B.D.,X.P., P.K.S., M.C.H.

Project administration: R.J.B.D., J.K.B., P.K.S., M.C.H.

Supervision: R.J.B.D., J.K.B., X.P., P.K.S., M.C.H.

Validation: R.J.B.D., J.K.B., X.P., P.K.S., M.C.H.

Visualization: R.J.B.D.

Writing: – original draft Writing: R.J.B.D

– review & editing : R.J.B.D., J.K.B., X.P., P.K.S., M.C.H.

