## Supplementary figures and images for "Fibroblast alignment is governed by stiffness interfaces in hydrogels"

### supp fig1

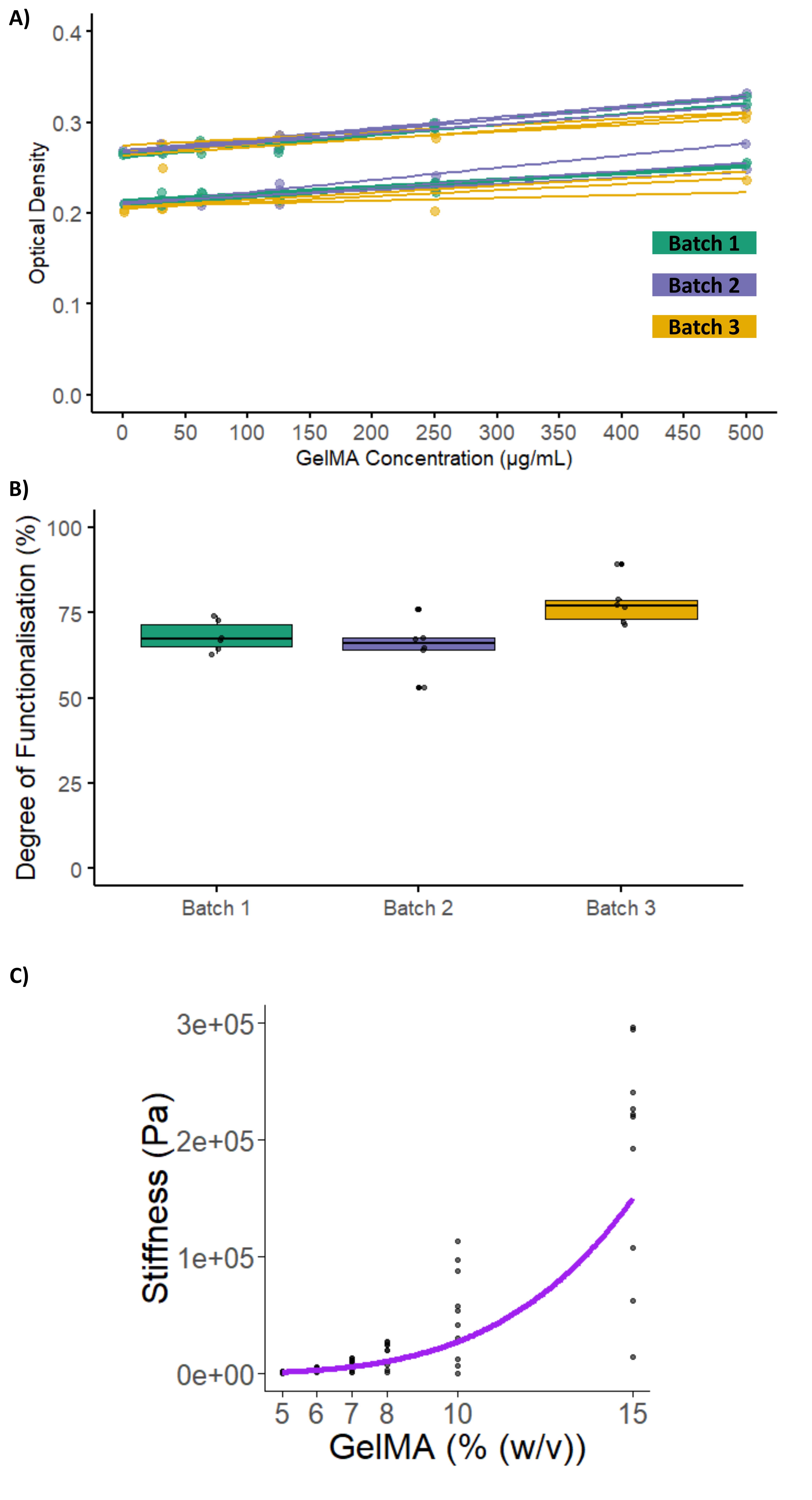

### supp fig2

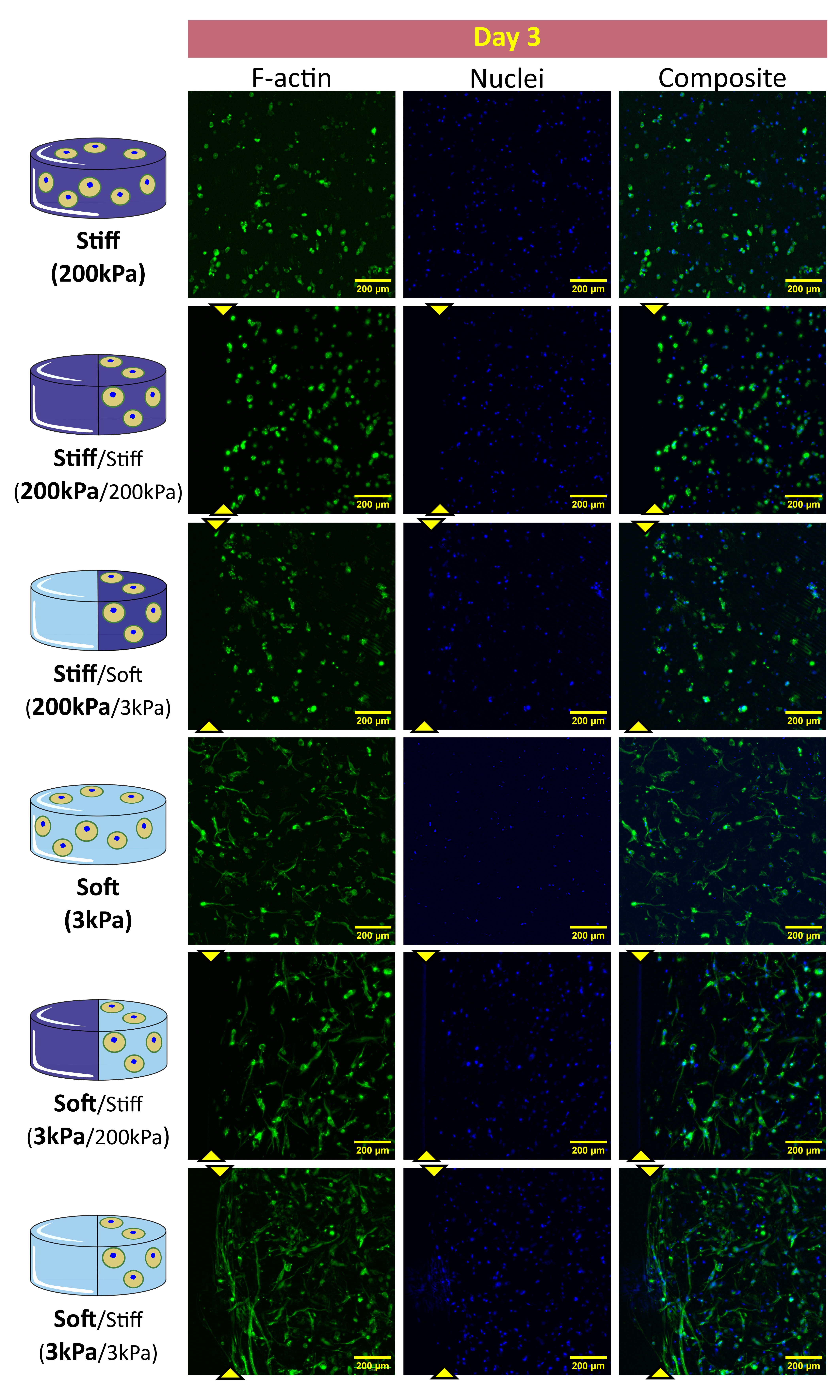

### supp fig3

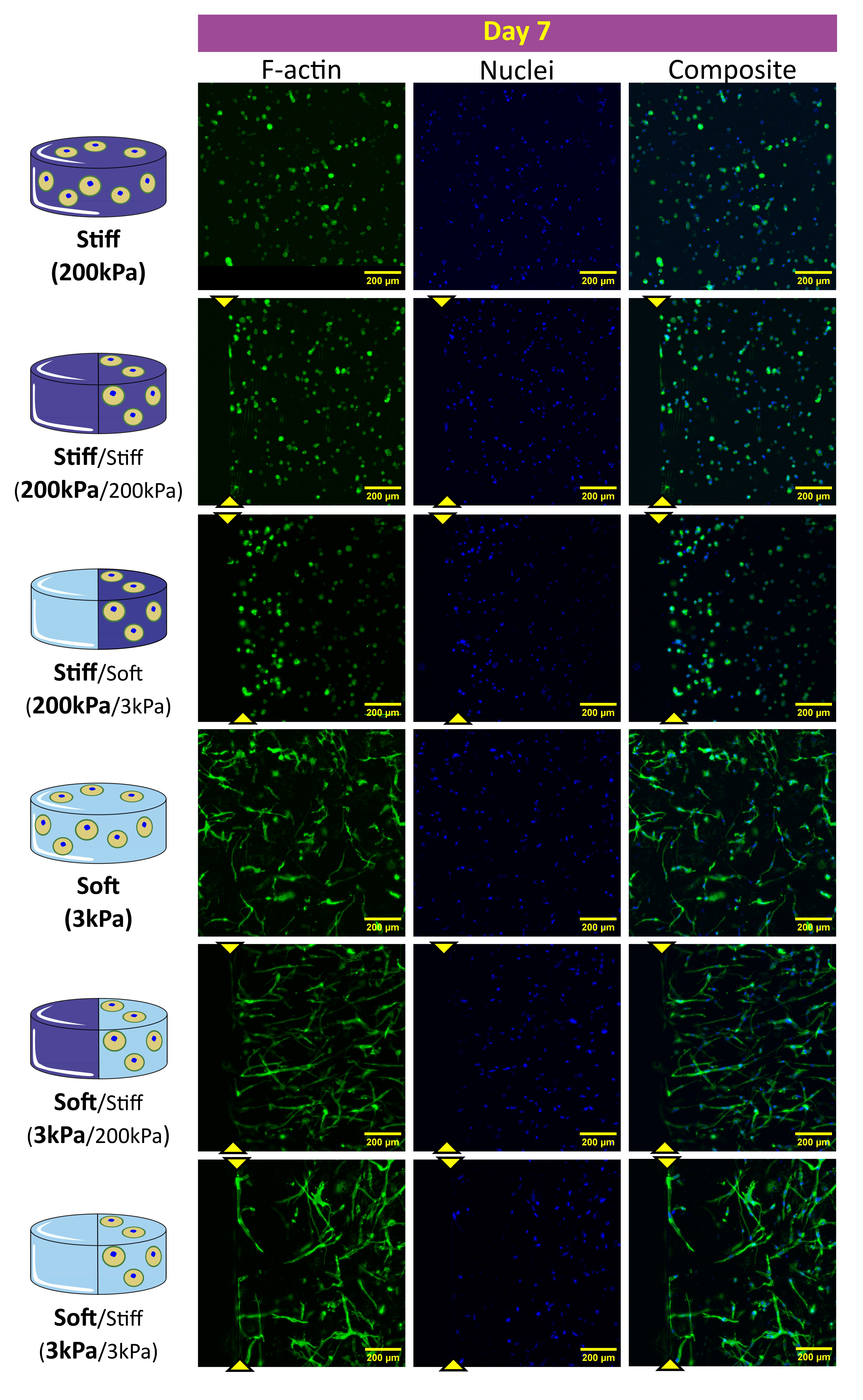

### supp fig4

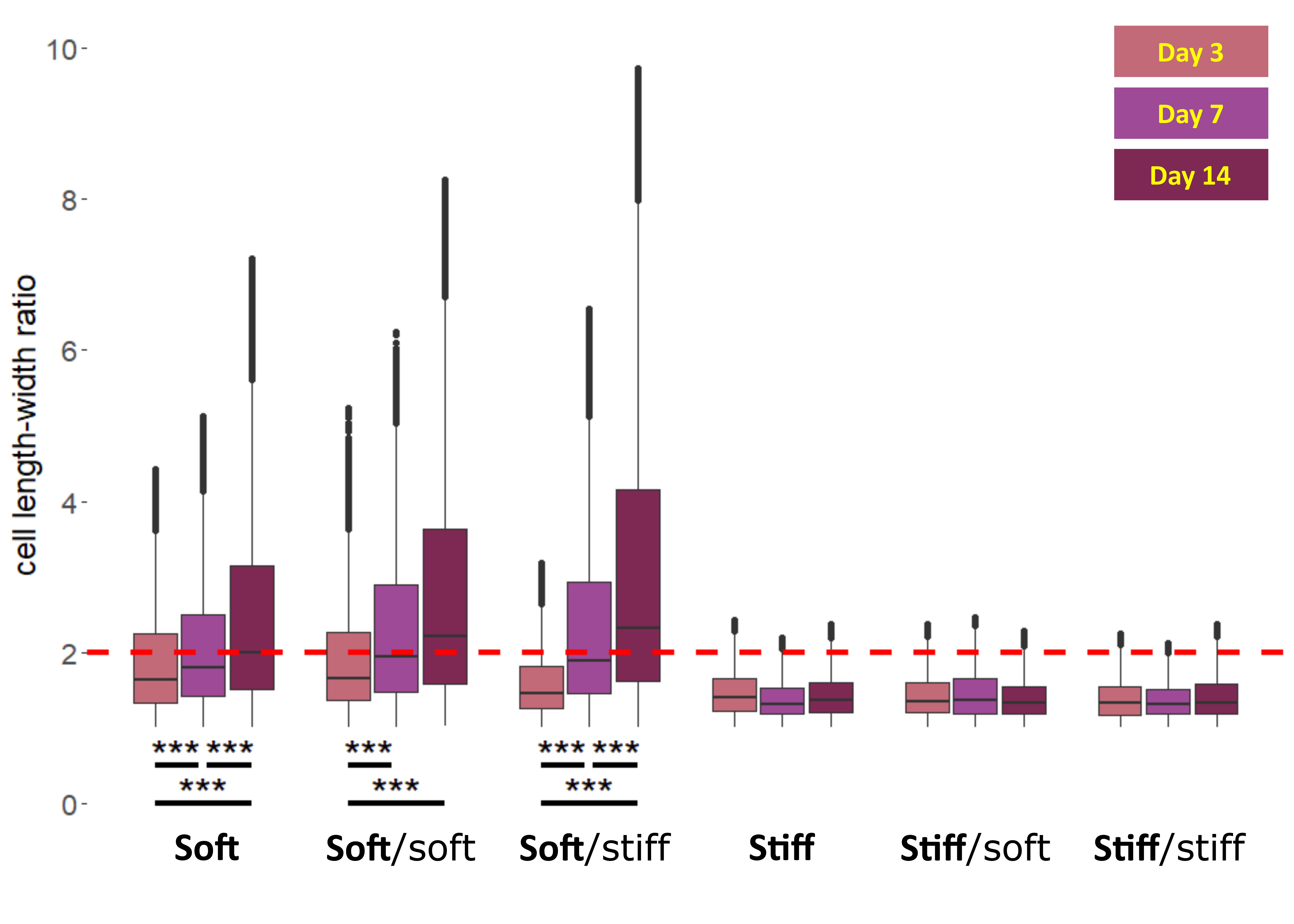

### supp fig5

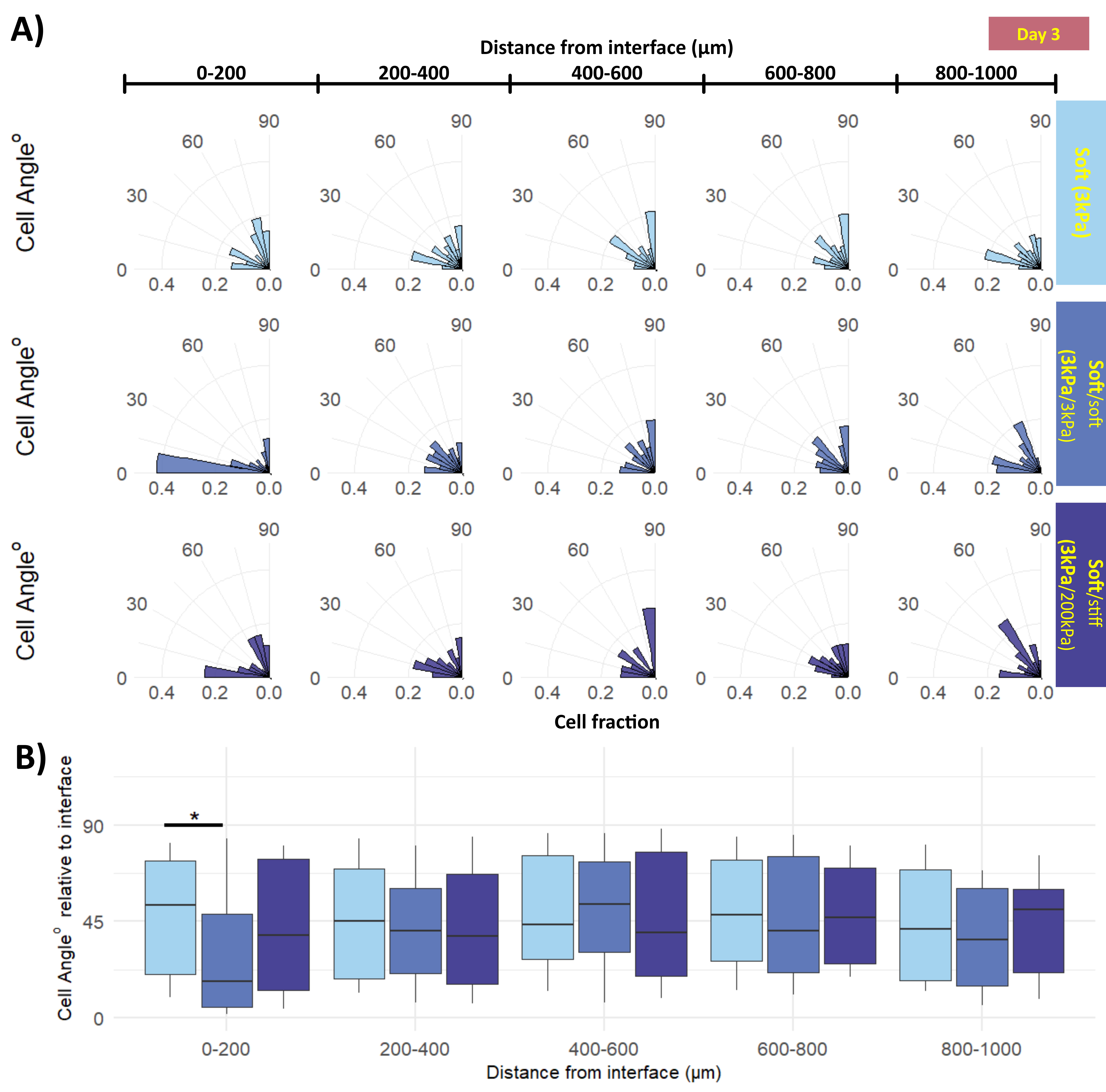

### supp fig6

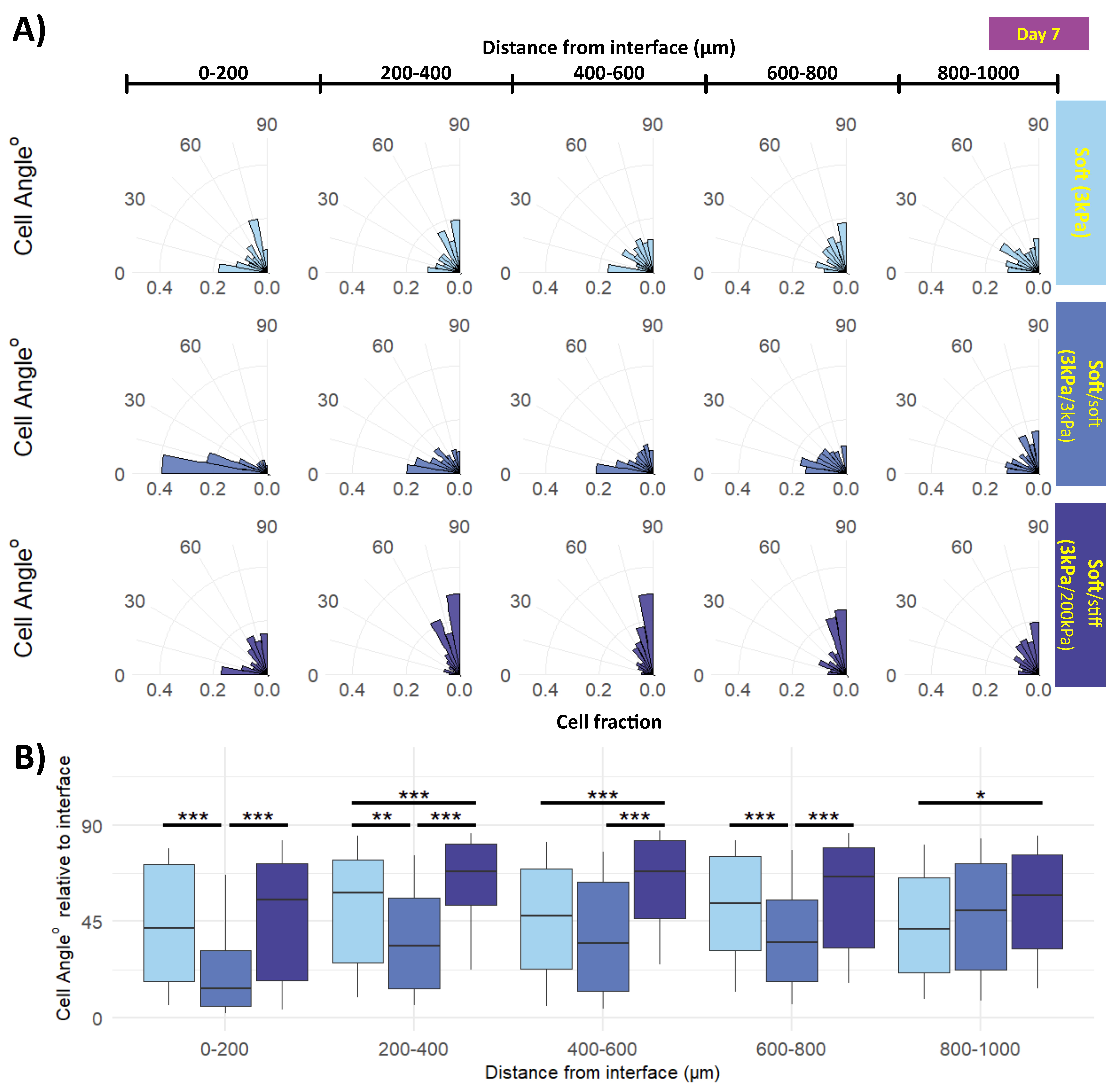
