## Supplementary material for "Fibroblast alignment is governed by stiffness interfaces in hydrogels": supp text

**Supplement 1**: **GelMA Characterization**
A) Shown are the six individual calibration curves for each batch (Batch 1 – green, Batch 2 – purple and Batch 3 – orange) to determine the degree of functionalization (DoF) with the 2,4,6-Trinitrobenzenesulfonic acid (TNBS) assay.
B) The average degree of functionalization of five hydrogels (black dots) per GelMA batch .
C) Calibration curve to determine the relation between GelMA concentration (%, w/v) and stiffness (kPa) (n > 8).

**Supplement 2:** **Stiffness transition driven fibroblast morphology and orientation.**
Fibroblast morphological and orientation differences after 3 days, influenced by the interface between GelMA hydrogels (indicated by the yellow arrows) composed of either soft (3kPa) or stiff (200kPa) hydrogel. The cell-containing hydrogel is indicated in **bold** font (1x10^6^ fibroblasts/ml). The cytoskeleton (Phalloidin) is shown in green and nuclei (DAPI) in blue. The scale bar represents 200 μm. Images are representative of three experimental repeats.

**Supplement 3:** **Stiffness transition driven fibroblast morphology and orientation.**
Fibroblast morphological and orientation differences after 14 days, influenced by the interface between GelMA hydrogels (indicated by the yellow arrows) composed of either soft (3kPa) or stiff (200kPa) hydrogel. The cell-containing hydrogel is indicated in **bold** font (1x10^6^ fibroblasts/ml). The cytoskeleton (Phalloidin) is shown in green and nuclei (DAPI) in blue. The scale bar represents 200 μm. Images are representative of three experimental repeats.

**Supplement 4:** **Hydrogel stiffness-dependent cell length-width ratio.**
Hydrogel stiffness-dependent cell length-width ratio as function of the culture time (d3, 7 and 14). **Bold** font indicates the cell-containing hydrogel with a seeding density of 1x10^6^ fibroblasts/ml. The median cell length-width ratio of a minimum of 750 fibroblasts per hydrogel is plotted, at least three individual hydrogels per interface group were measured. Cells with a length-width ratio over 2 (dash redline) were considered elongated. ANOVA with Tukey post-hoc analysis was used to assess differences between the groups. (* p<0.05, ** p<0.01 and *** p<0.001).

**Supplement 5: Stiffness interfaces direct fibroblast alignment.**
A) The distribution in cell angle of fibroblasts (°) relative to the interface within GelMA hydrogel at day 3 of culture. The fraction of cell angles is plotted for **soft** (light blue), **soft**/soft (lavender), and **soft**/stiff (purple).
B) Median cell angle relative to the interface derived from at least 30 fibroblasts per distance in at day 14 of culture for **soft** (light blue), **soft**/soft (lavender), and **soft**/stiff (purple). Statistical differences were assessed using Kruskal-Wallis with Dunn’s post hoc test (*p < 0.05, **p < 0.01, ***p < 0.001).
From left to right, the distance from the interface is given in bin ranges of 200 μm, up to 1000 μm distance from the interface.

**Supplement 6: Stiffness interfaces direct fibroblast alignment.**
A) The distribution in cell angle of fibroblasts (°) relative to the interface within GelMA hydrogel at day 7 of culture. The fraction of cell angles is plotted for **soft** (light blue), **soft**/soft (lavender), and **soft**/stiff (purple).
B) Median cell angle relative to the interface derived from at least 30 fibroblasts per distance in at day 14 of culture for **soft** (light blue), **soft**/soft (lavender), and **soft**/stiff (purple). Statistical differences were assessed using Kruskal-Wallis with Dunn’s post hoc test (*p < 0.05, **p < 0.01, ***p < 0.001).
From left to right, the distance from the interface is given in bin ranges of 200 μm, up to 1000 μm distance from the interface.
