## Supplementary material for "Fibroblast alignment is governed by stiffness interfaces in hydrogels": table1

**Table 1: Fibroblast-mediated modulation of hydrogel stiffness**

|  | Incubation Time  (Days) | Stiffness (kPa) | | p-value |
| --- | --- | --- | --- | --- |
|  |  | 0 cell/ml | 1x10^6^ cell/ml |  |
| Soft | 0 | 2.9 (IQR: 2.4-3.3) | 4.7 (IQR: 3.3-6.9) | 0.01 |
|  | 3 | 2.4 (IQR: 2.2-2.6) | 3.7 (IQR: 2.9-4.7) | 0.03 |
|  | 7 | 2.0 (IQR: 1.7-2.4) | 3.4 (IQR: 2.4-5.7) | 0.03 |
|  | 14 | 2.1 (IQR: 1.8-2.4) | 2.3 (IQR: 2.0-3.0) | 0.23 |
| Stiff | 0 | 229 (IQR: 201-260) | 138 (IQR: 124-154) | 0.01 |
|  | 3 | 194 (IQR: 185-202) | 100 (IQR: 98-171) | 0.04 |
|  | 7 | 190 (IQR: 172-306) | 109 (IQR: 56-134) | 0.01 |
|  | 14 | 173 (IQR: 163-177) | 107 (IQR: 63-128) | 0.01 |

GelMA hydrogel stiffness was measured by LLCT in the presence and absence of fibroblasts (1x10^6^ fibroblasts/ml) after culture for up to 14 days. The stiffness (kPa) is shown as median of five replicates per time point with interquartile range (IQR: 25-75%). Kruskal-Wallis with Dunn’s post-hoc analysis was used to assess differences between groups, represented in p-values.
