## Supplementary material for "Fibroblast alignment is governed by stiffness interfaces in hydrogels": table2

**Table 2: Fibroblast-mediated modulation of hydrogel stress relaxation**

|  | Incubation Time  (Days) | Stress Relaxation (%) | | p-value |
| --- | --- | --- | --- | --- |
|  |  | 0 cell/ml | 1x10^6^ cell/ml |  |
| Soft | 0 | 5.9 (IQR: 4.5-7.8) | 5.6 (IQR: 4.7-5.8) | 0.45 |
|  | 3 | 5.9 (IQR: 5.8-6.1) | 6.5 (IQR: 5.9-8.0) | 0.35 |
|  | 7 | 6.9 (IQR: 6.6-7.2) | 6.3 (IQR: 4.0-9.0) | 0.67 |
|  | 14 | 6.9 (IQR: 6.3-7.3) | 8.1 (IQR: 7.7-8.2) | 0.02 |
| Stiff | 0 | 6.1 (IQR: 4.2-7.4) | 3.9 (IQR: 3.8-4.2) | 0.12 |
|  | 3 | 8.5 (IQR: 6.6-13.9) | 3.4 (IQR: 3.0-3.4) | 0.01 |
|  | 7 | 7.2 (IQR: 5.6-8.6) | 3.5 (IQR: 3.1-4.1) | 0.01 |
|  | 14 | 7.6 (IQR: 5.7-9.3) | 3.1 (IQR: 3.0-3.7) | 0.01 |

GelMA hydrogel stress relaxation was measured by LLCT in presence and absence of fibroblasts presence (1x10^6^ fibroblasts/ml) after culture for up to 14 days. The stress relaxation (%) is shown as median of five replicates per time point with interquartile range (IQR: 25-75%). Kruskal-Wallis with Dunn’s post-hoc analysis was used to assess differences between groups, represented in p-values.
